# A dataset of differentiable biologically-derived single neuron models

**DOI:** 10.64898/2025.12.02.691895

**Authors:** Calvin Yeung, Zhixin Lu, Kris Ganjam, Stefan Mihalas

## Abstract

Biological neural networks contain diverse cell types with heterogeneous electrophysiological properties. Artificial neural networks (ANNs) model computational aspects of biology but use homogeneous neurons, limiting realism. A key obstacle to building bio-inspired ANNs is the absence of a database of neuron models that both fit biological data and remain differentiable for gradient-based learning. We present such a database: over 1,000 differentiable single-neuron firing-rate models with accompanying PyTorch code for integration with machine learning. The models belong to the linear-nonlinear (LN) class, which, for ease of reference, we term Generalized Firing Rate (GFR) neurons. Each GFR neuron uses input and firing-rate history filters across multiple timescales, followed by a nonlinearity to generate firing rates. Models are fit to patch-clamp recordings from mouse and human central nervous system slices. Parameter clustering reflects electrophysiological diversity and partially aligns with transgenic lines. This resource will enable development of bio-inspired ANNs built from biologically grounded, differentiable single-neuron models.

**Highlights:**

- Resource of over 1,000 differentiable single-neuron models fit to mouse and human patch-clamp recordings
- Open source PyTorch code and reproducible pipelines support model fitting, evaluation, and network integration
- Fitted model parameters capture electrophysiological diversity, with parameter clusters partially aligning to inhibitory/excitatory types and Cre driver lines

## 1. Introduction

The intersection of neuroscience and machine learning has increasingly revealed computational parallels between biological and artificial neural networks [1]. While machine learning has made significant improvements by drawing inspiration from the neural mechanisms of the brain, typical neuron models used are incredibly simple, utilizing a transfer function on a weighted sum of inputs. Such models overlook the intricate dynamics and diversity found in biological neurons, which are comprised of hundreds of distinct types [2].

Biologically-informed single neuron models could be used in building artificial neural networks designed to understand the mechanisms of computations in biological circuits. One possible reason for the paucity of such models is a lack of proper tools. This is what we plan to alleviate with this resource: we introduce a dataset of over 1,000 computationally efficient and fully differentiable neuronal models which have been individually fit to the patch-clamp recordings of mouse and human cortical neurons[3]. The code is implemented in PyTorch[4], which is one of the most used tools in machine learning. Crucially, the single neuron models are designed to remain backpropagation-trainable when integrated into network architectures, allowing integration into larger deep learning architectures while maintaining the capacity to train individual neuron parameters. The GFR model is quite interpretable, with the main mechanisms being driven by two types of currents: one driven by the history of the input, and one by the history of the neuron’s activity.

There is a vast literature on single neuron models which we will only briefly summarize. For a comprehensive but older summary please see [5] or a more recent but not as complete review [6]. Mammalian cortical neurons have extensive dendrites that can result in non-linear summation of inputs, with individual neurons being as complex as 5-8 layers deep neural networks [7]. Following dendritic integration, multiple types of ion channels [8] at the soma transform the input current into a set of action potentials. The set of models that we construct focuses on the somatic transformation from input currents to neuronal activity. The reason for this is twofold: (1) In the large-scale measurements of synaptic transmission between mammalian cortical neurons, post-synaptic potentials or currents are typically measured at the soma [9, 10], and so the filtering done by the dendrites is already included in the measurements. (2) Large-scale measurements of the single neuron in vitro physiology are performed with somatic current injection [11].

Indeed, there is interest in including biophysical details in simulating the dynamics of neural networks. For example, in [12], they constructed networks consisting of biophysically realistic neurons and used backpropagation to optimize not only the connections but also the biophysical parameters of individual neurons. However, in that study, the biophysical parameters of the individual neurons are obtained through task-driven optimization, as no database of individual fly neuron models was available.

The two closest studies that constructed model databases on the same dataset we use are [13] and [14]. However, these models are intended for the neuroscience community and are not differentiable. While the measure of explained variance is not identical, the fitted GFR models achieve a similar explained variance ratio of 72% with a time bin of 20ms compared to GLIF’s 70-78% and biophysically detailed models 65-69% with a 10ms Gaussian smoothing (which gives a ± 1*σ* interval of 20ms). More generally, searching the database of 838 single neuron model studies hosted on ModelDB [15], we did not find a single dataset of differentiable models fit to biological data.

In contrast to GLIF models [14, 16] and LSTM/GRU [17, 18], which apply multiple nonlinearities to the history of various parameters, and dendritic models, which introduce multiple nonlinearities to the inputs, the GFR model is a linear-nonlinear model with a linear filter on the input and activity history followed by a nonlinearity in the activation function, making it easier to optimize via backpropagation. While LSTM and GRU are differentiable, they lack biological realism. As we will show in the next section, the GFR shares structural similarities with state-space models (SSM) [19, 20, 21]. Thus, our contributions are as follows:

1. We curate and release a large dataset of parameters for differentiable single-neuron models fit to patch-clamp recorded mouse and human cortical neurons.
2. We provide a fully reproducible training and evaluation pipeline so others can refit or extend the models.
3. We establish baseline results and usage examples to demonstrate the dataset’s utility.

## 2. The GFR Neuron Model

To capture dynamics in a computationally efficient yet biologically realistic manner, we consider linear-nonlinear firing rate models that describe the neuron’s firing rate *f* in response to input currents *I*. We make the simplifying assumption that the firing rate is determined by a non-linear activation function *g*(·) with history dependence, which can be described by a set of filters on the input and activity. Specifically,

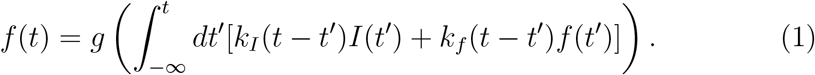

Since we generally expect a neuron’s temporal dependence on input stimulus and firing rate to decay over time, we choose the kernels *k*_*I*_ and *k*_*f*_ to be weighted sums of *n* decaying exponential functions, i.e.,

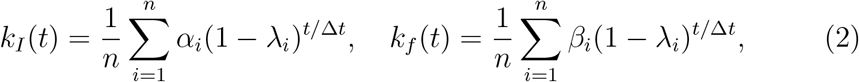

where *λ*_*i*_ ≤ 0 are the exponential decay rates, *α*_*i*_ and *β*_*i*_ are exponential weights, and Δ*t* > 0 is an arbitrary time constant to ensure that the argument of the exponential function is dimensionless. The factor 1/*n* is added for normalization. For practical purposes, we discretize the time-integral of Eq. 1 into bin sizes of Δ*t* milliseconds, giving us the approximation

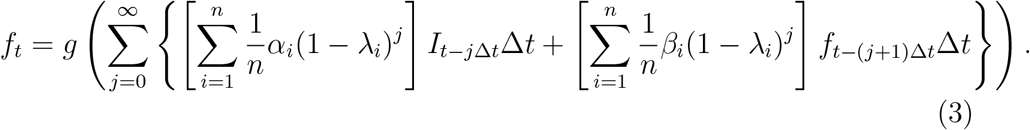

The exponential kernels in Eq. 3 require explicit summation over historical data. Here, we avoid such a direct summation over history by utilizing latent variables with a leaky-integration mechanism of different decay rates, 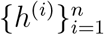, achieving flexible temporal convolutions.

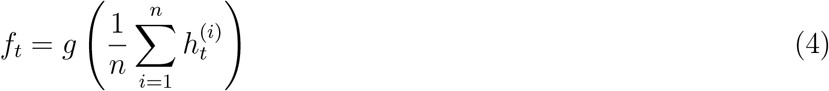

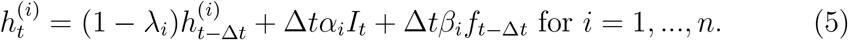

We set the first decay rate *λ*_1_ = 1, corresponding to a component in the GFR model that processes only the current input *I*_*t*_ and the previous firing rate *f*_*t*−1_. Figure 1 provides a schematic of the GFR model.

**Figure 1.**
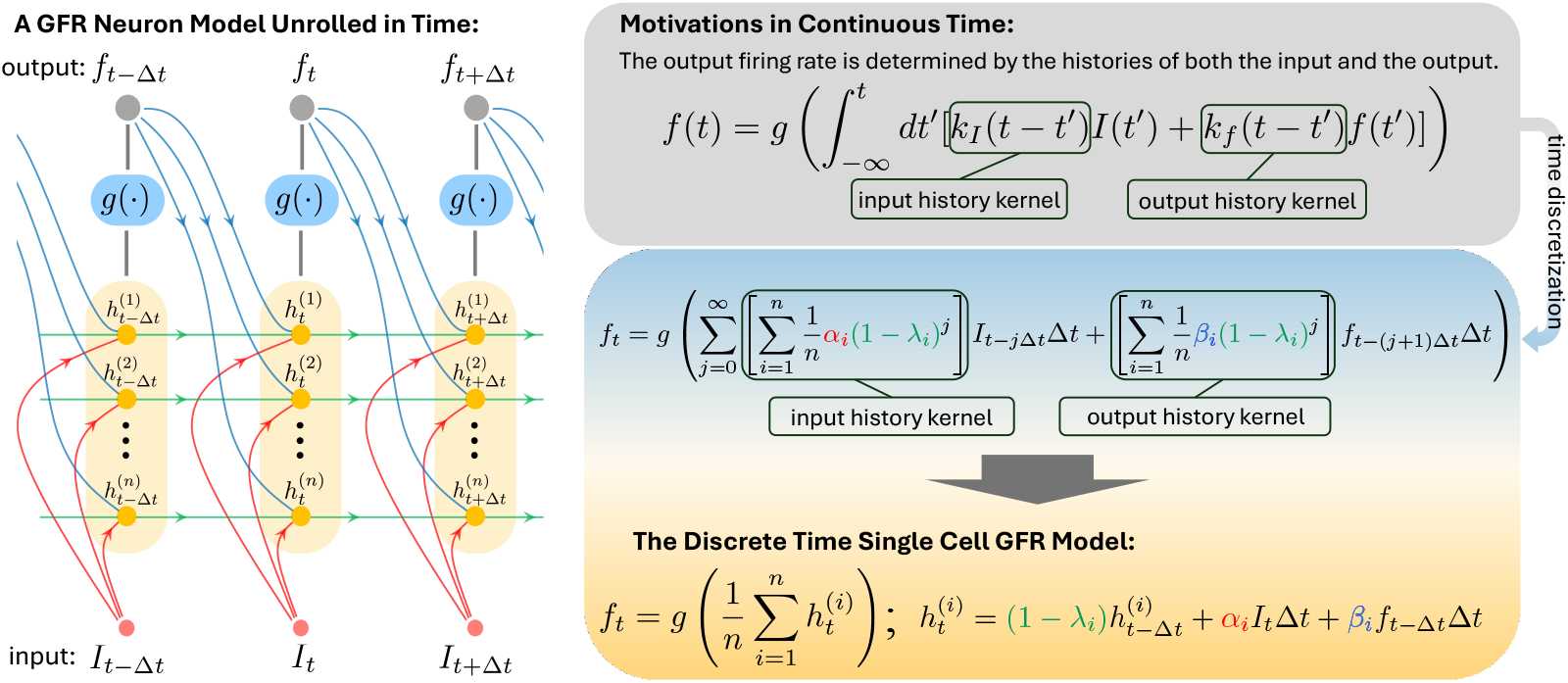
A schematic of a single GFR neuron unrolled in time. The GFR neuron takes input current *I*_*t*_ and outputs firing rate *f*_*t*_. The GFR model aggregates the input and output history in hidden variables 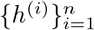, performing temporal convolutions over input and output history.

We use an activation function *g* defined as

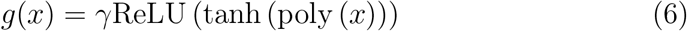

is a polynomial of degree *d*. For ease of optimization, we write 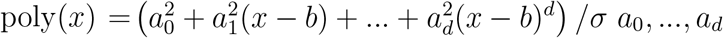 are trainable parameters. We square *a*_*i*_ to ensure the coefficients are non-negative. We pre-compute *γ*, the maximum firing rate of the neuron, *b*, the firing threshold, and *σ*, the maximum experimental current. *γ* and *σ* are fixed during training.

In comparison to SSMs that can model complex dynamical responses using linear dynamics, the GFR model also employs linear ordinary differential (or difference) equations for its hidden states 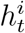, which represent different ion current dynamics. However, unlike the interacting states *x*_*i*_(*t*) in SSMs, GFR’s states evolve independently with unique decay timescales. While SSMs typically produce outputs through linear functions, the GFR model uses a nonlinear activation function tailored to neuronal firing dynamics. This function not only captures the nonlinear relationship between ion concentrations and firing rates but also includes a tunable maximum firing rate, making the GFR model more biophysically interpretable.

## 3. Fitting GFR Parameters to Electrophysiology Data

### 3.1. The Electrophysiology Dataset

To obtain a dataset of GFR neurons, we fit the GFR parameters to the Allen Cell Types Electrophysiology Database [11, 3] which consists of patch clamp recordings of individual neurons from the mouse and human cortex labeled by fluorescent proteins with an expression pattern determined by a Cre-based driver. A variety of driver lines were used to sample broadly across the cortical circuit as well as to target specific excitatory and inhibitory populations. For each cell, the dataset records spike times alongside voltage in response to various types of stimulus currents. Stimulus types include Long Square, Short Square, Noise 1, Noise 2, Ramp, etc. Among most stimulus types, there are multiple amplitudes of currents. For example, a Long Square stimulus may consist of a constant current for 1 second with an amplitude of 30 pA.

### 3.2. Preprocessing the Electrophysiology Data

#### Preprocessing for Kernel Parameter Optimization

We convert spike data into firing rate data via bin-averaging on both the current and spike data. In particular, we use bin sizes Δ*t* = 10, 20, 50, 100 milliseconds. Moreover, before bin-averaging, we truncate the start of the sequence where zero input current is present so that the onsets of the stimuli are aligned with the beginning of time bins.

#### Preprocessing for Activation Parameter Optimization

To fit the activation function, we focus only on Long Square stimuli. From the Long Square sweeps, we produce current-firing rate pairs, where the current is the amplitude of each Long Square stimulus, and the firing rate is the firing rate computed at the first time bin with non-zero current.

### 3.3. Optimization of the GFR Activation Function

#### Computing Maximum Current and Firing Rate

To fit the activation function *g*(·) defined in Eq. 6, we first compute the maximum firing rate *γ* by taking the minimum difference between spike times, converting it into milliseconds, and then taking the reciprocal. We also compute the maximum experimental current *σ* by taking the maximum bin-averaged current over all stimuli with activation bin size Δ*t*′. During the optimization, we fix *γ* and *σ*.

#### Initialization of Activation Parameters

We initialize the firing threshold *b* to be the smallest current that induces a non-zero firing rate in the preprocessed dataset for the activation function. We initialize polynomial coefficients *a*_2_, …, *a*_*d*_ ~ *N* (0, *ϵ*) for some small *ϵ*. Coefficients *a*_0_, *a*_1_ are computed with an approximate linear function passing through the current-firing rate pair with the largest firing rate and the pair with the smallest current with a non-zero firing rate.

#### Optimization Details

To optimize the parameters, we use the Adam optimizer [22], minimizing the Poisson negative log-likelihood between the predicted number of spikes per time bin and ground truth. We optimize activation parameters for bin sizes Δ*t*′ = 20, 100, using *d* = 1 for Δ*t*′ = 20 and *d* = 3 for Δ*t*′ = 100. We did not use an activation bin size of 10 as the sparsity of spikes led to poor fits for many cells. We considered that a fine-grained (20 ms) and coarse-grained (100 ms) fit was sufficient for the estimation of the transfer function.

### 3.4. Optimization of Exponential Kernel Parameters

#### Choosing Decay Rates

We use exponential kernels with timescales *τ* chosen from a list of 10, 20, 50, 100, 200, 500, 1000, and 2000 milliseconds. If we use a bin size Δ*t*, we only use kernel timescales *τ* ≥ Δ*t* among the list. To convert a timescale *τ* to decay rate *λ* used in Eq. 5, we use the equivalence *λ* = 1 − *e*^−Δ*t*/*τ*^. Alongside the decay coefficients computed using the time bins specified above, we add a decay coefficient *λ*_1_ = 1 to model instantaneous temporal behavior.

#### Initialization of Kernel Parameters

The GFR neuron is initialized as a memoryless neuron. Specifically, for the current kernel coefficients *α*_*i*_, we set *α*_1_ = *n* and the remaining *α*_*i*_ = 0 for *i* = 2, …, *n*. We initialize all firing rate coefficients *β*_*i*_ = 0 for *i* = 1, …, *n*. This way, the initial behavior of the GFR neurons matches the behavior of the computing units in artificial neural networks, where the neuron output is solely determined by the instantaneous input via an activation function. We use activation parameters previously computed as described above and keep them fixed.

#### Data Splitting and Batching

We place Long Square, Ramp, and Noise 1 sweeps into the train dataset and Noise 2 sweeps into the test dataset. We batch the bin-averaged data (with bin size Δ*t*) according to stimulus type, zero-padding as necessary.

#### Optimization Details

We optimize the model over bin sizes Δ*t* = 10, 20, 50, 100 milliseconds with Adam and use the average Poisson negative log-likelihood loss over each batch and sequence as the objective. We apply *L*_0_ regularization on the kernel parameters, with regularization strengths *C* ∈ {1, 0.5, 0.1, 0.05, 0.01, 0.005, 0.001, 0}. We initialize hidden variables 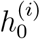 and firing rate *f*_0_ to zero at the start of each sweep. For each bin size Δ*t*, we use an activation function with bin size Δ*t*′ ≥ Δ*t*. For each Δ*t*, Δ*t*′ pair, we perform model selection using the explained variance ratio on Noise 1.

#### Toy use example

In appendix Appendix C we provide an example of how GFR neurons can be used in a network to solve a toy machine learning problem, sequential MNIST [23, 24], in which a network has to classify handwritten digits, but the image information available to the network is provided sequentially.

#### Code availability

We include a zip of code that can be used to reproduce the results in the supplementary. The code, as well as the parameter sets, is also freely available at https://github.com/AllenInstitute/GRNN/.

## 4. Results

### 4.1. GFR Neurons Fit to the Cell Types Dataset

Below are examples of the fitted GFR models along with a quantitative evaluation of their fitting quality across all the trained GFR neurons.

#### Example Fitted GFR Neurons

We visualize the fitting results for four neurons from the Allen Cell Types Electrophysiology Database [11]: (a) a Pvalb cell (567901050); (b) a Vip cell (560814288); (c) an Rbp4 cell (486132712); and (d) an Ntsr1 cell (488674910). Cell IDs are specified in parentheses. Figure 2 plots the activation functions (Δ*t*′ = 100 milliseconds) of the chosen neurons, while Figure Appendix B.1 plots the corresponding learned kernel parameters and kernels.

**Figure 2.**
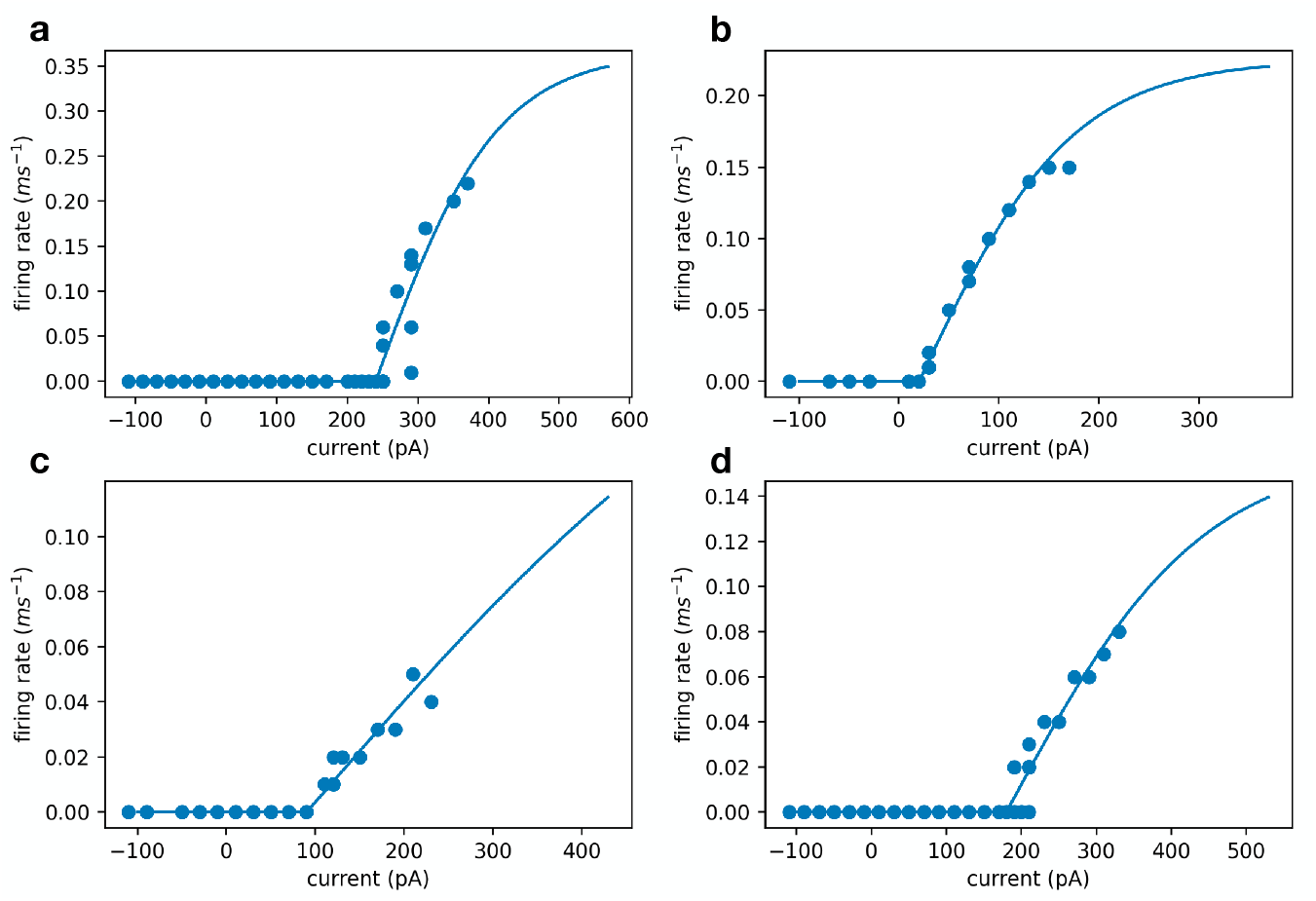
Plot of learned activation functions with Δ*t*′ = 100 milliseconds. **a**.The activation function of a Pvalb cell (567901050), exhibits a very high max firing rate. **b**. The activation function of a Vip cell (560814288), which has a very small firing threshold. **c**. The activation function of an Rbp4 cell (486132712). **d**. The activation function of an Ntsr1 cell (488674910).

To visualize the firing rate behavior of the chosen neurons to different stimulus types, we also plot the input stimulus along with closed-loop firing rate predictions for four cells in the dataset for GFR models trained on Δ*t* = 20 milliseconds in Figure 3. For each cell, we plot the Long Square, Ramp, and Noise 1 stimulus types. The predicted firing rate approximates the average ground truth firing rate at any given time point.

**Figure 3.**
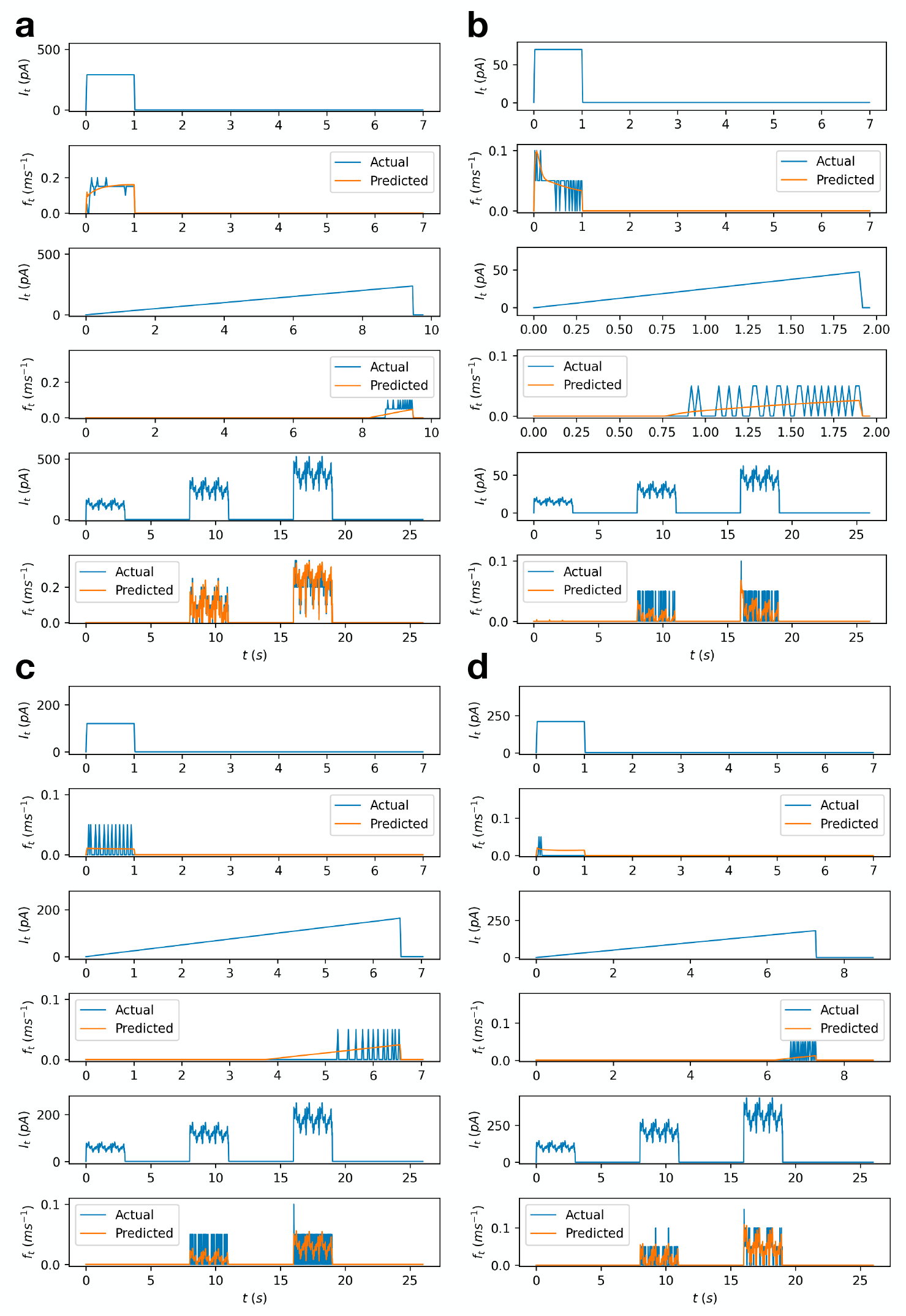
Closed-loop GFR firing rate predictions for four cells over Long Square, Ramp, and Noise 1 stimuli for Δ*t* = 20 milliseconds. **a, b, c**, and **d** correspond to the cells in Figure 2.

#### Model Evaluation Over All Neurons

To evaluate the fitting quality of all the GFR neurons, we compute the explained variance ratio for all cells on the Noise 2 data as in [16, 14], but with bin-averaging as described in Section 3.2. Figure 4 visualizes the distribution of explained variance ratios and reports the median explained variance ratio for each bin size over all cells in the dataset that have Noise 1 and 2 sweeps present in the data.

**Figure 4.**
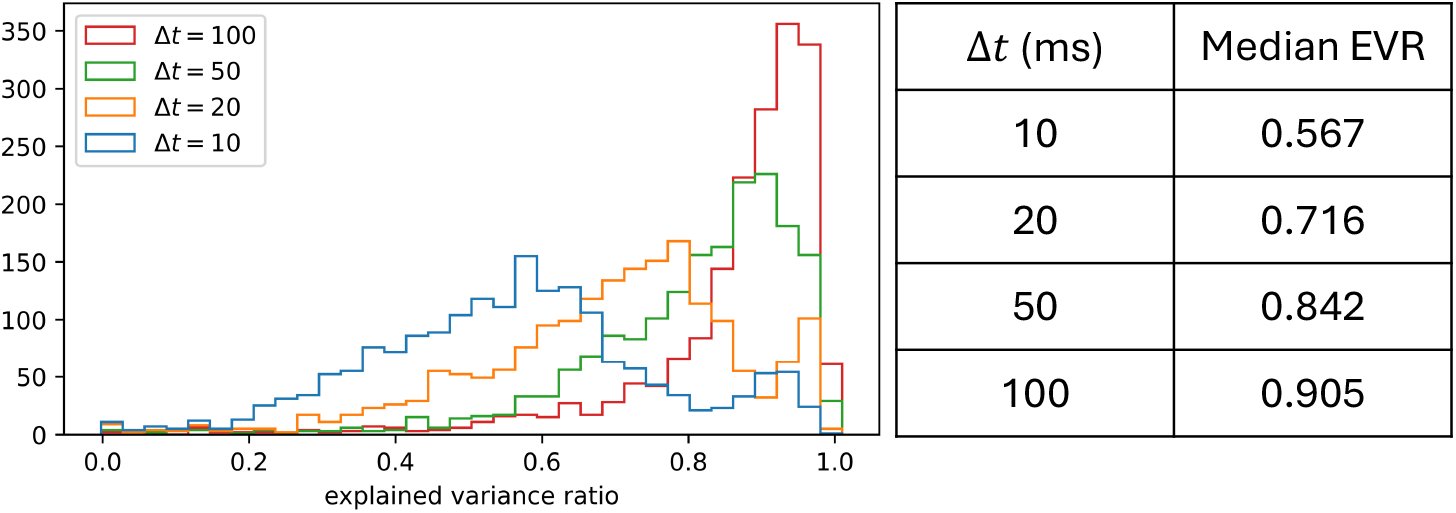
**Left**: A histogram of the computed explained variance ratio for GFR models on testing data (Noise 2) with bin sizes of 10, 20, 50, and 100 milliseconds over the entire dataset. **Right**: The median explained variance ratio for the different bin sizes.

#### Selection Criteria for GFR Dataset

The GFR dataset we present consists of trained GFR model parameters for different configurations of bin sizes and activation bin sizes. We only include models that pass certain criteria, namely: (1) the data includes both Noise 1 and Noise 2 sweeps; (2) the validation explained variance ratio on Noise 1 (which is the training dataset) is greater than 0.5; and (3) the training loss is less than 0.45. Table Appendix C.2 lists the number of cells satisfying the above criteria for different configurations.

### 4.2. Parameter Clustering

To prove the utility of this dataset, we aim to link clusters in parameter space of the models to transgenic lines. We perform ward hierarchical clustering on the learned parameters for all models with an explained variance ratio greater than 0.5 on Noise 2. We use a bin size and activation bin size of 20 milliseconds. As preprocessing, we replace features *a*_0_, *a*_1_, *b, σ* with *c*_0_, *c*_1_, where 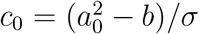 and 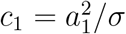, to remove redundancy. Note that we square the polynomial coefficients *a*_0_, *a*_1_ here since we do the same in the implementation to ensure the coefficients are non-negative. In addition, we perform z-score normalization for each parameter.

#### Clusters Reflect Biological Features

For learned GFR models of each cell type, we visualize its cluster membership as shown in Figure 5A. In general, we see a trend where cluster membership correlates with the inhibitory and excitatory nature of the cells. We also find that parvalbumin cells are very distinct as expected. We obtain an adjusted Rand index of 0.104 when the number of clusters is 11.

**Figure 5.**
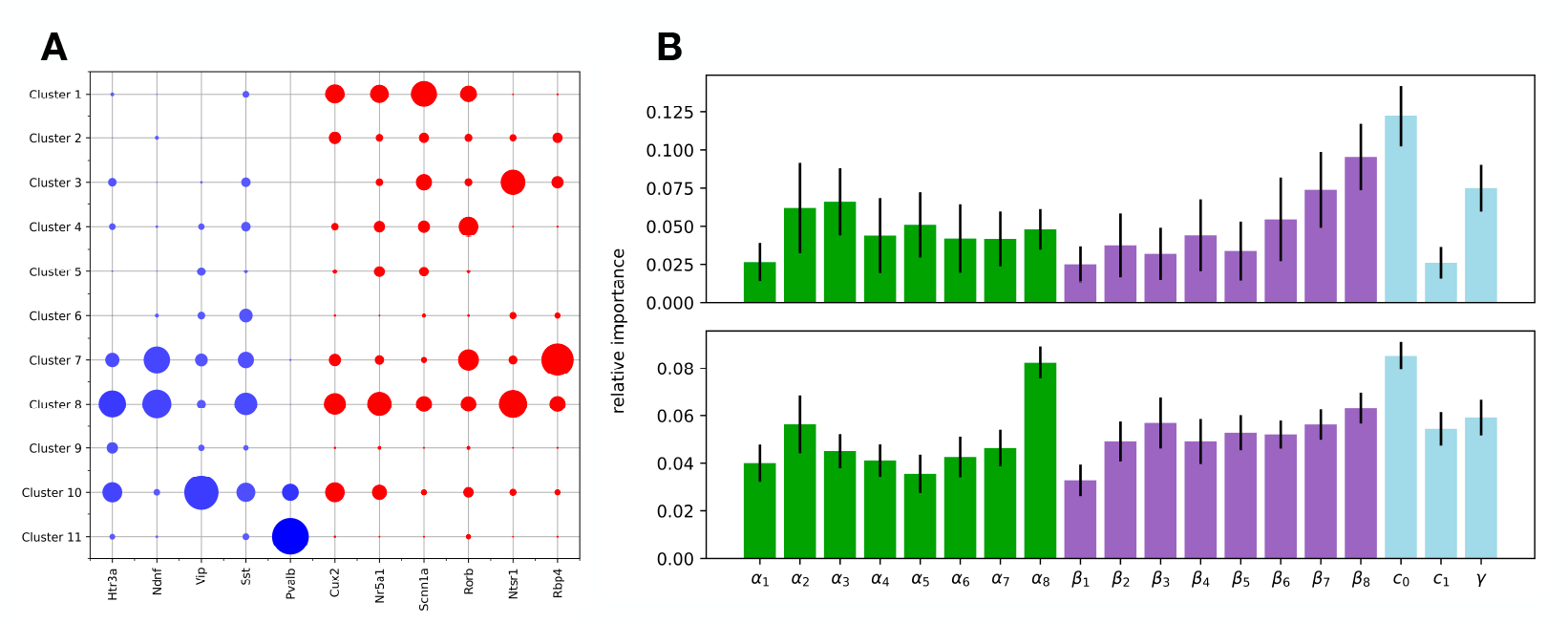
**A**. Cluster visualization by cre-line. We perform hierarchical clustering on learned neuron parameters (Δ*t*, Δ*t*′ = 20). The size of the circle denotes the number of cells of a given cell type in a given cluster, normalized by the total number of cells of that cell type. Red circles are inhibitory and blue circles are excitatory. **B**. Average importance weights computed using a GLM for clustering over multiple optimization runs. Error bars indicate standard deviation. Top: importance weights with respect to predicting inhibitory vs excitatory cells. Bottom: importance weights with respect to predicting cre-line. The green, purple, and light blue bars correspond to current history kernel, firing rate history kernel, and activation parameters respectively.

#### Computing Relative Importance

To compute the relative importance of each feature, we train a generalized linear model **y** = softmax(**Mx** + **b**) to classify parameters by cre-line. The resulting importance weights are computed as 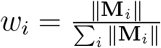, where **M**_*i*_ is the *i*-th column of **M** and ∥ · ∥ is some norm. In this case, we use the *L*_2_ norm. Figure 5B visualizes the learned importance weights for each of the features averaged over multiple optimization runs. We find that when predicting whether cells are inhibitory or excitatory, activation parameters are more important. On the other hand, when predicting cre-line, while activation parameters are still important, kernel parameters are also more predictive.

## 5. Conclusion and Discussion

Our biologically-derived, differentiable single-cell models can provide a unique tool to study computations by neural networks, particularly in contrast to traditional artificial neural networks (ANN), enriching machine learning (ML) with biological realism.

### Distinction from Traditional ANN Units

Traditional artificial neural networks predominantly utilize simple activation functions such as sigmoid, ReLU, or tanh, which process inputs without an inherent mechanism to incorporate complex temporal dependencies [25]. In stark contrast, our single-cell model integrates historical information through its structure. This integration is achieved using linear temporal filters that model the decay and interaction of past inputs and neuronal states, reflecting a more dynamic and temporally aware computational framework. This capability allows the model to mimic more closely the temporal dynamics inherent in biological neurons, where the state history and past inputs significantly influence current behavior. Such features are especially critical in tasks involving time series predictions, decision-making processes, and dynamic environment interactions, where historical context enhances performance and decision accuracy.

#### Towards Better Tools for Neuroscience Models

Our models are directly derived from high-quality electrophysiological recordings of cortical neurons, embodying the complex and rich dynamics observed in biological systems. This biological grounding introduces realism into neural network simulations, offering researchers tools to study fundamental computational differences between biological neurons and conventional ML models. Our database of fully differentiable single-neuron models allows other researchers to embed biologically realistic cells into large-scale networks and examine their impact on learning, robustness, and circuit-level dynamics.

#### Towards Biologically-Informed Machine Learning

The introduction of our dataset and modeling approach represents a step towards a more biologically-informed kind of machine learning. This shift could pave the way for more sophisticated learning algorithms that not only achieve high performance but also exhibit properties such as generalization, fault tolerance, and energy efficiency—characteristics inherent in biological systems but often lacking in artificial constructs.

#### Limitations

The experimental dataset underlying the model database is not uniformly sampled from the cortex, with a bias in mice for primary visual cortex and in humans for medial temporal lobe. The experiments are performed in the absence of neuromodulator, which will affect the single neuron input-output relations. The experiments are performed with somatic injections, and as such the dendritic nonlinearities are not captured. For mice, we looked at the relation between neuron models and originating transgenic lines, which each contain multiple transcriptomic cell types. The model is constructed to have a unique nonlinearity.

## Appendix

**A. GFR Dataset Details**

**Table Appendix A.1:**
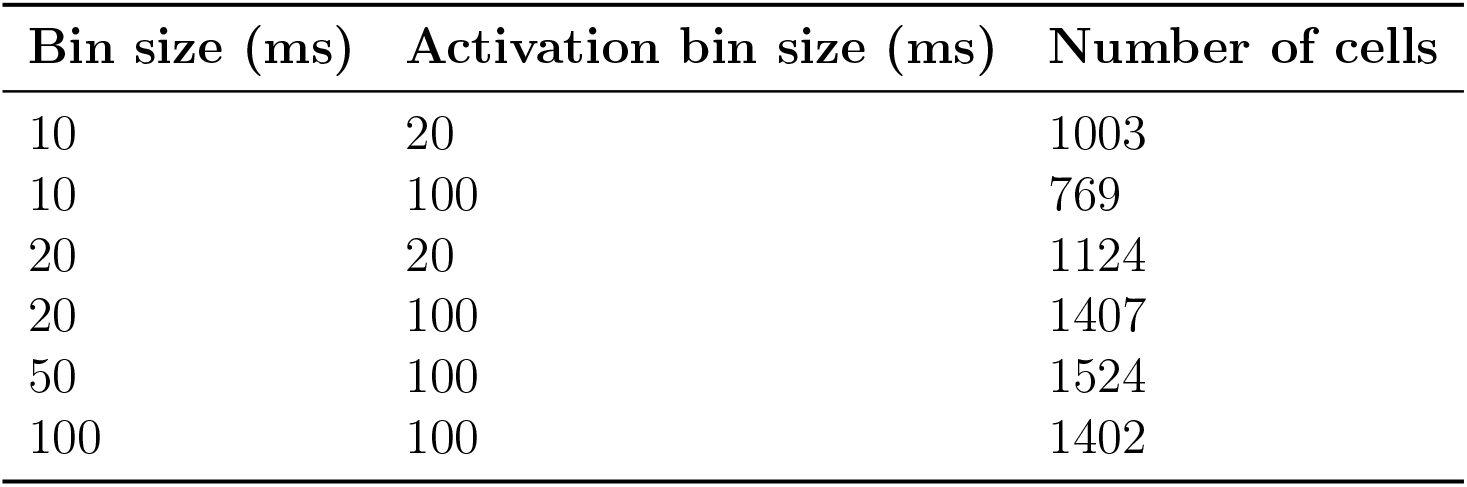
Size of dataset which pass all the inclusion criteria for different bin size and activation bin size configurations.

| Bin size (ms) | Activation bin size (ms) | Number of cells |
| --- | --- | --- |
| 10 | 20 | 1003 |
| 10 | 100 | 769 |
| 20 | 20 | 1124 |
| 20 | 100 | 1407 |
| 50 | 100 | 1524 |
| 100 | 100 | 1402 |

## Appendix

**B. GFR Parameters for Example Neurons**

**Figure Appendix B.1:**
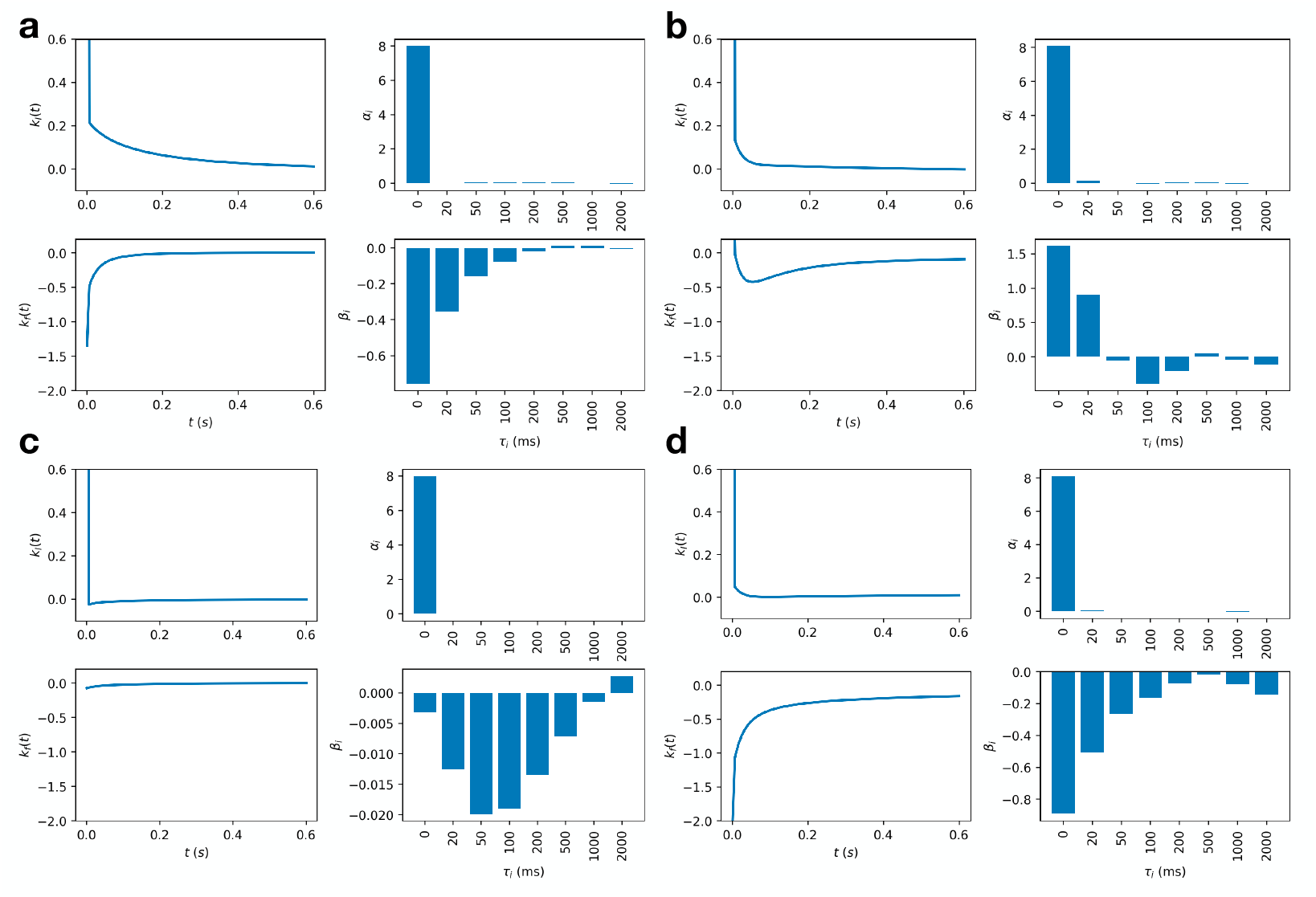
Plot of learned kernel parameters and kernel shape. **a, b, c**, and **d** correspond to the cells in Figure 2.

## Appendix

**C. Sequential MNIST**

## Appendix C.1. Solving the L-MNIST Task by an RNN of GFR Neurons

To demonstrate the versatility and computational power of our GFR neuron models, we construct a recurrent neural network (RNN) to solve the L-MNIST task [24] using a hidden layer of 64 GFR neurons. L-MNIST, or line-by-line MNIST, converts the original MNIST dataset into a sequential dataset by converting each 28 × 28 image into a sequence of 28 rows or columns. Given the task’s inherently discrete nature in time, we set Δ*t* = 1. Figure Appendix C.1 illustrates the sequential MNIST task and the general recurrent neural network schematic.

Since biological neurons did not evolve to solve L-MNIST, we first did not use GFR parameters fit to biological data (see appendix Appendix C.2 for GFR-RNN performance using biological parameters). Instead, we initialized each GFR neuron as in Section 3.4 and trained them using backpropagation from scratch. In addition, we use a fixed activation function with *γ, σ* = 1, *b* = 0. We also use a polynomial with degree *d* = 1 such that poly(*x*) = *x*.

**Figure Appendix B.2:**
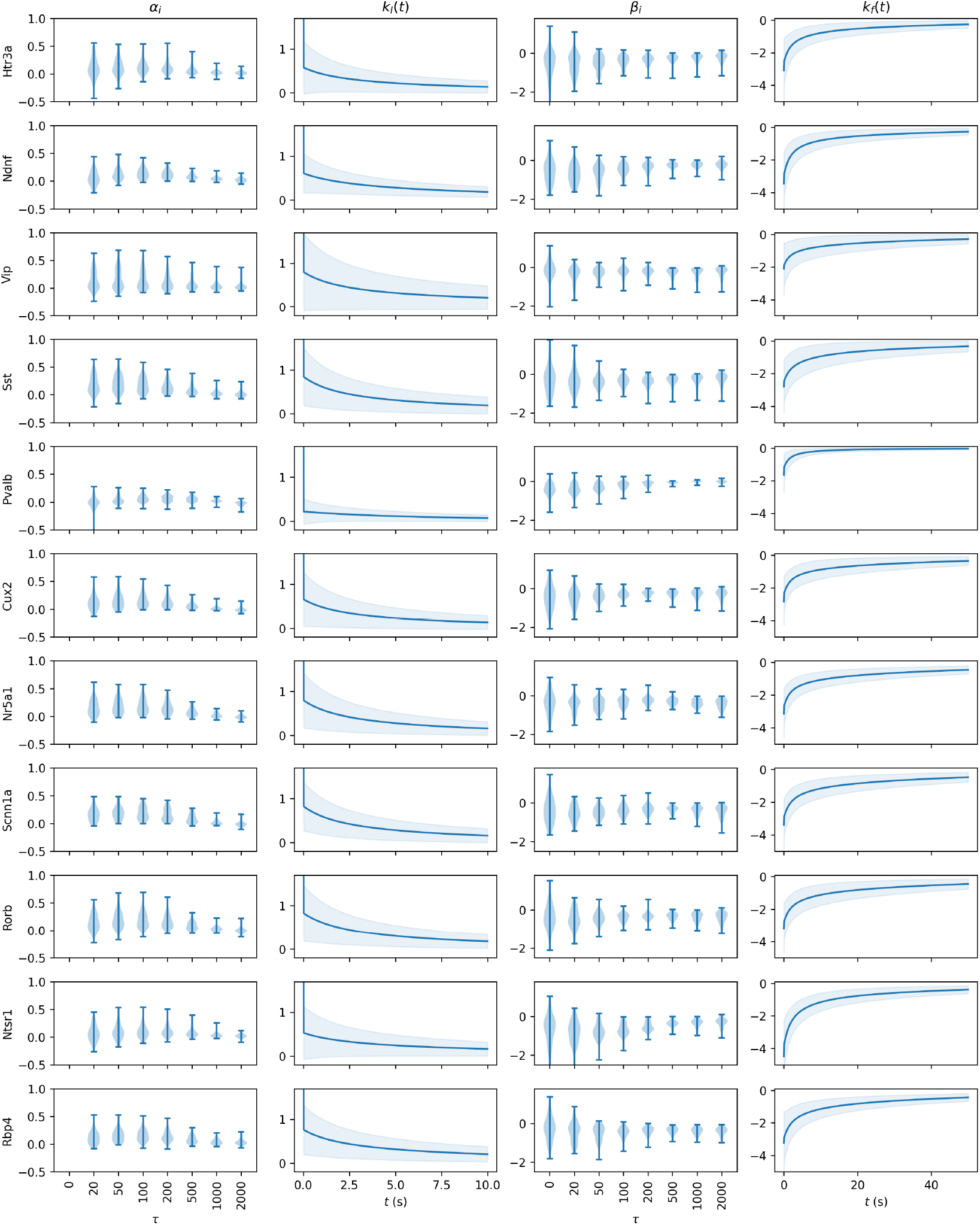
Plot of kernel parameter and kernel distribution by cre-line. For the kernel plots, the blue line is the average kernel, while the shaded region indicates the standard deviation.

**Figure Appendix B.3:**
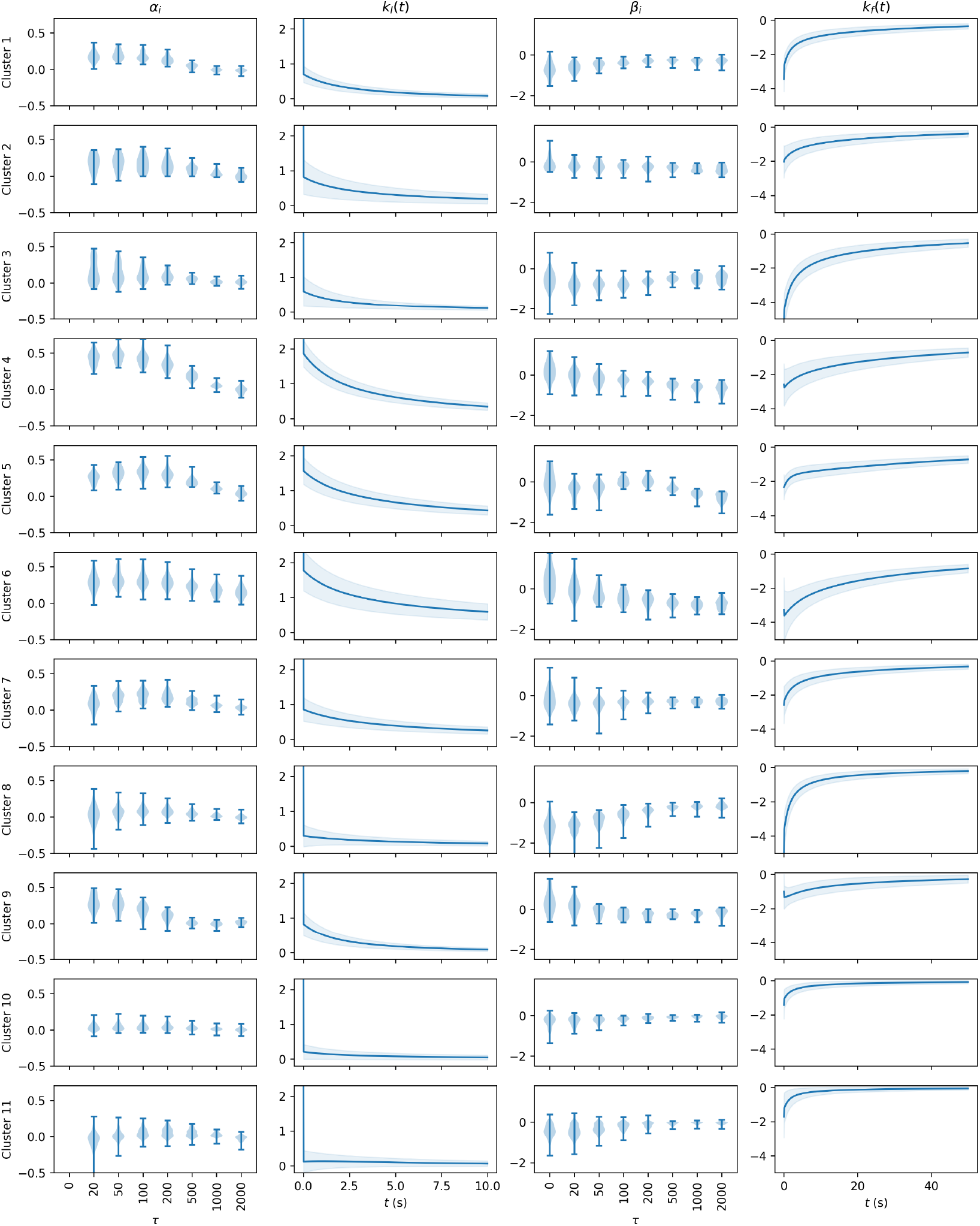
Plot of kernel parameter and kernel distribution by found clusters. For the kernel plots, the blue line is the average kernel, while the shaded region indicates the standard deviation.

**Figure Appendix B.4:**
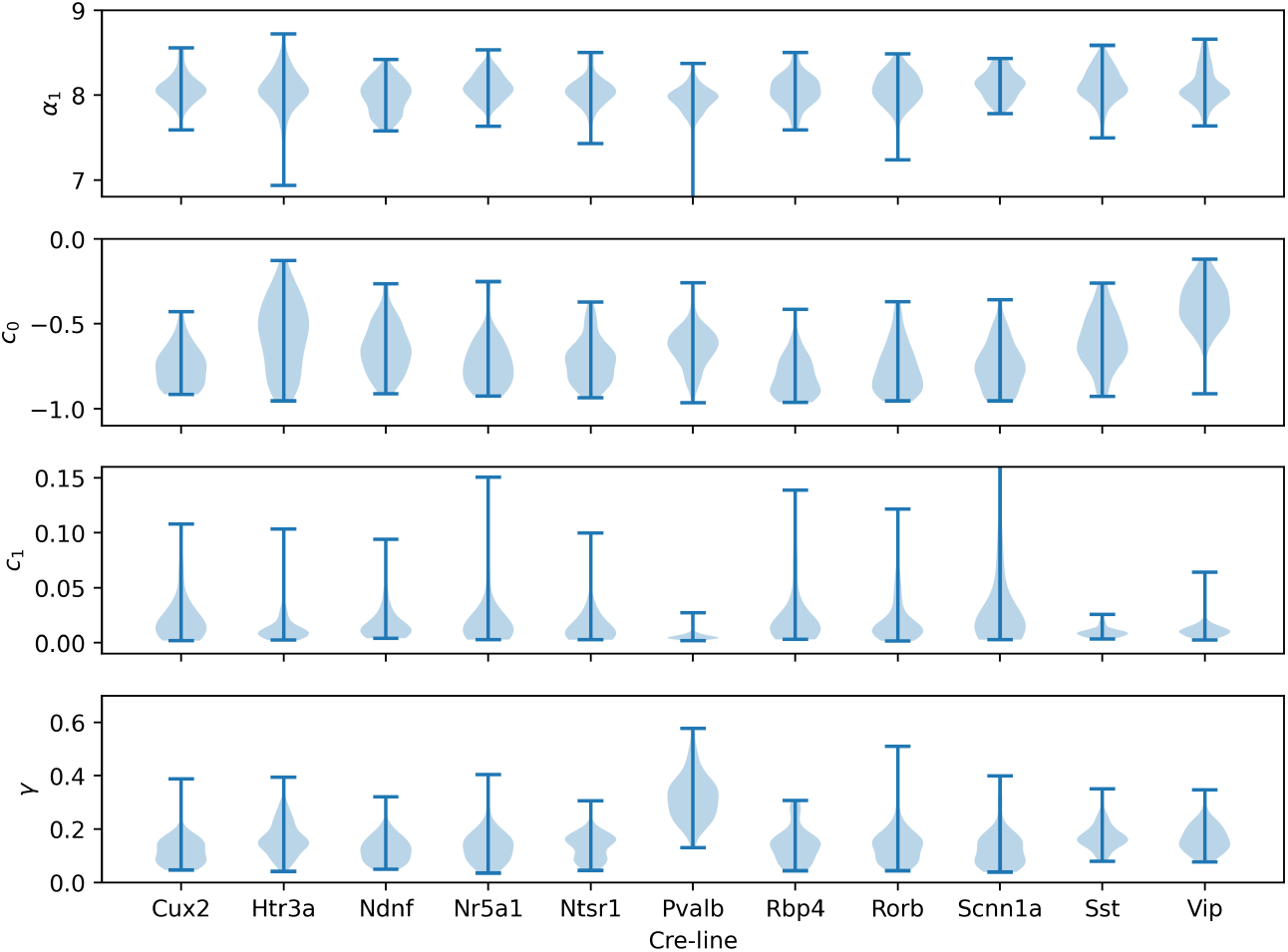
Plot of parameters *α*_1_, *c*_0_, *c*_1_, *γ* by cre-line.

**Figure Appendix B.5:**
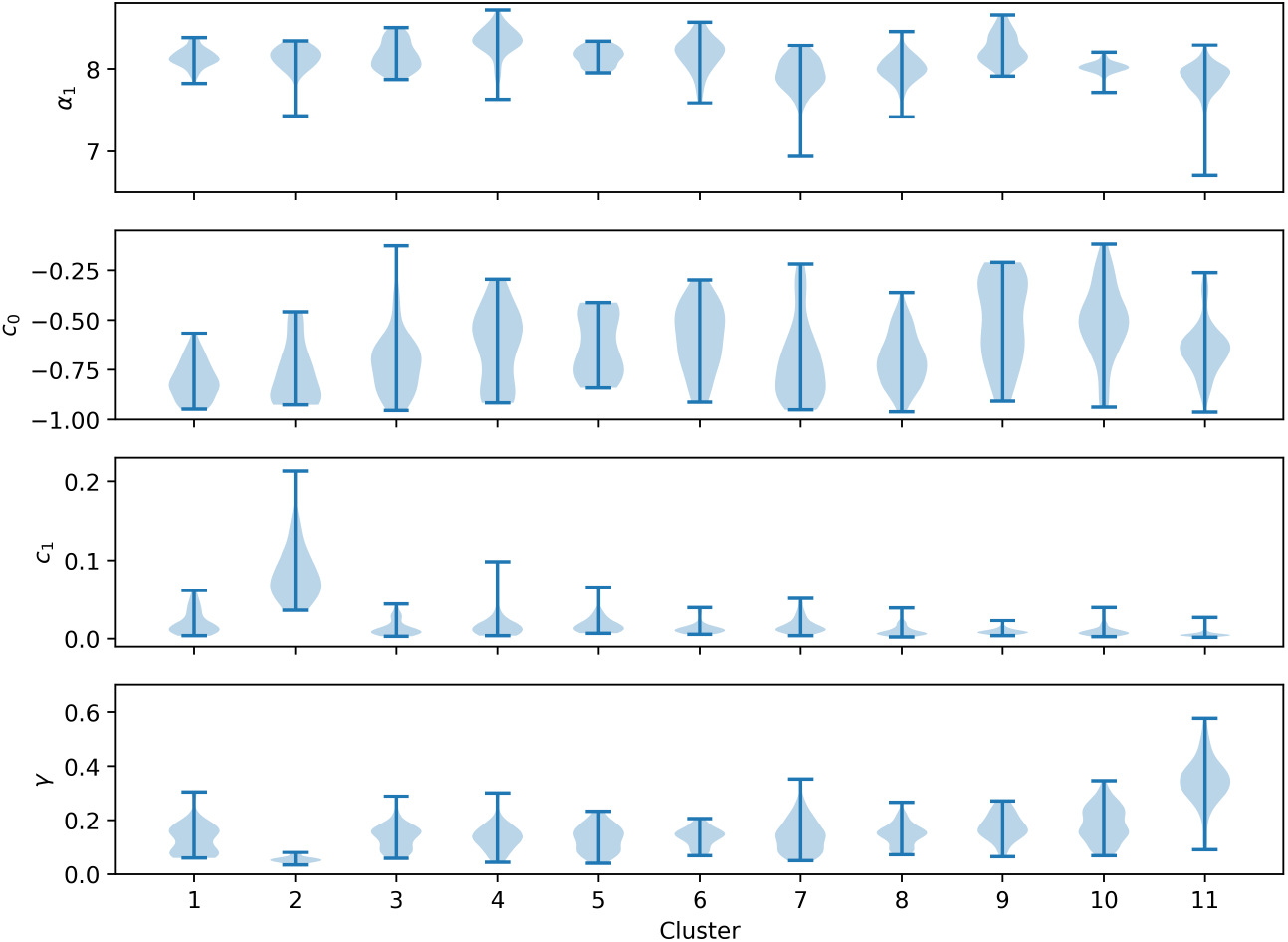
Plot of parameters *α*_1_, *c*_0_, *c*_1_, *γ* by found clusters.

**Figure Appendix C.1:**
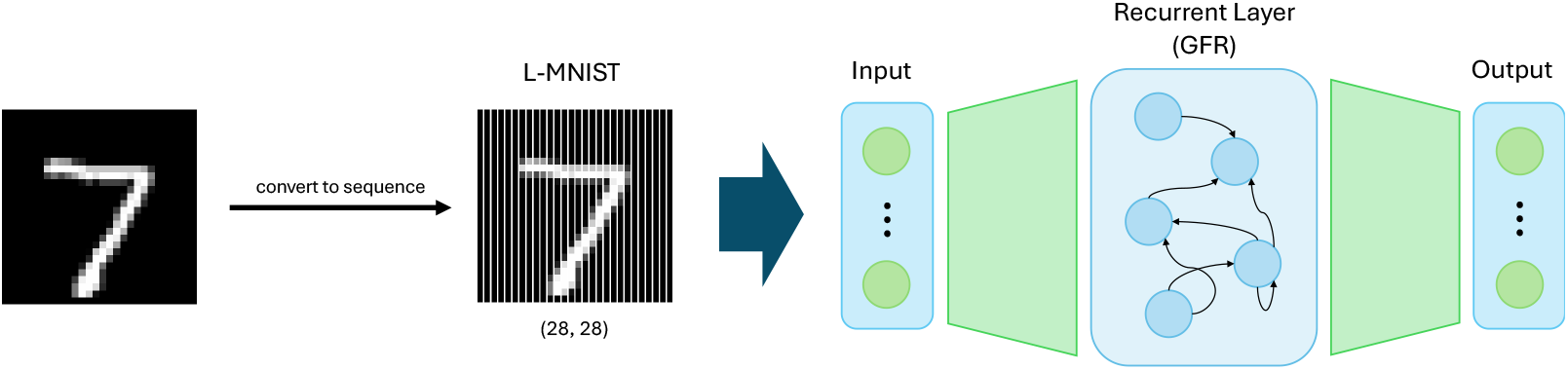
The sequential MNIST pipeline. An image from the MNIST dataset is converted into a sequence and then passed sequentially into a recurrent neural network with GFR neurons. Finally, a prediction is made.

### The GFR-RNN Architecture

The GFR-RNN consists of one input layer, one hidden layer, and one output layer. Let *N* be the input-layer dimension, *D* the hidden-layer dimension, and *M* the output-layer dimension. We denote the input vector at time step *t* by **x**_*t*_ ∈ ℝ^*N*^, the GFR neurons (i.e. hidden layer) activities by **z**_*t*_ ∈ ℝ^*D*^, and the GFR-RNN’s output vector by **y**_*t*_ ∈ ℝ^*M*^. The *D* GFR neurons in the hidden layer receive input currents from both the input layer and the hidden layer, represented by a *D*-dimensional vector,

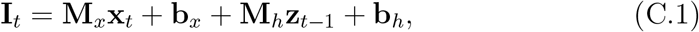

where its *i*-th entry, [**I**_*t*_]_*i*_, corresponds to the *i*-th neuron’s input current *I*_*t*_ in Eq. 5. The *i*-th entry of **z**_*t*_ is the firing rate of the *i*-th GFR neuron, defined in Eq. 4. The GFR-RNN’s output is given by

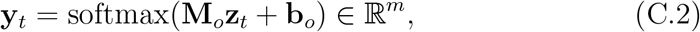

where **M**_*x*_, **M**_*h*_, **M**_*o*_, **b**_*x*_, **b**_*h*_, **b**_*o*_, together with the *α*’s and *β*’s in Eq. 5 for each of the GFR neurons, are the GFR-RNN’s learnable parameters.

### Training Details

For a given sequence of length *T*, we make a prediction by taking the maximum entry in **y**_*T* +1_, where the input to the RNN after time *T* is the zero vector **0** ∈ ℝ^*n*^. During training, we backpropagate based on the cross-entropy between the ground truth and **y**_*T* +1_. We use Adam to optimize the GFR-RNN model with a learning rate of 1e-3 and train over 300 epochs.

**Figure Appendix C.2:**
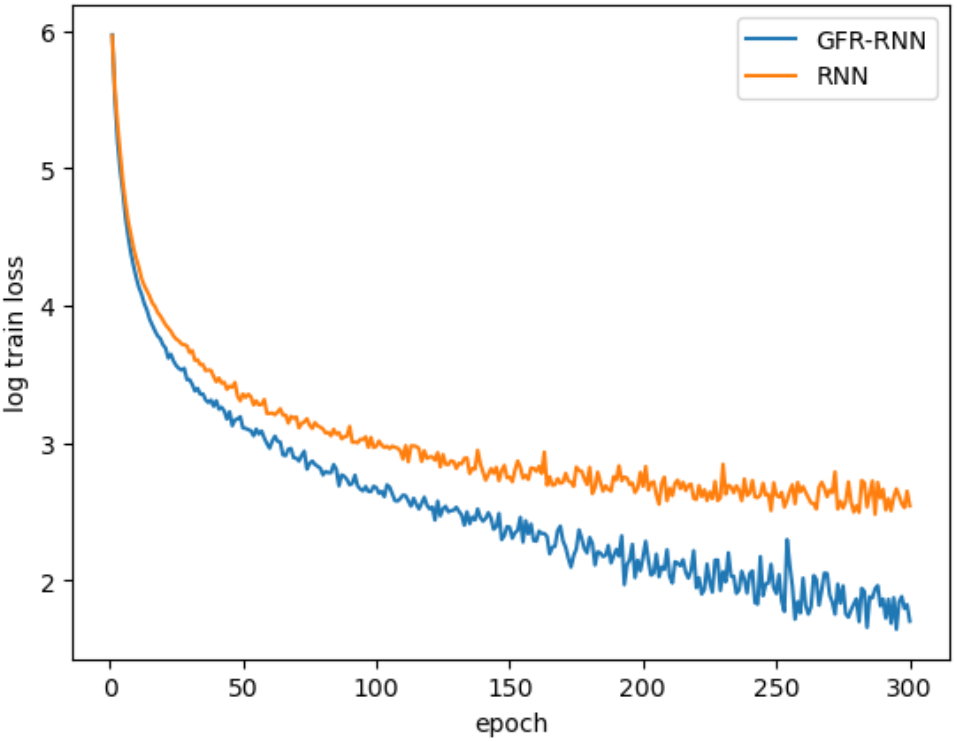
Log train loss for both GFR-RNN and RNN models over 300 training epochs.

### Comparing to the Standard RNN

In this GFR-RNN, each GFR neuron is equipped with kernels of timescales *τ* = 1, 2, 5 alongside an instantaneous kernel *τ* = 0. For the activation function, we set a maximum firing rate *γ* of 1 and used a simple polynomial poly(*x*) = *x*. Table Appendix C.1 reports the test accuracy of the trained models and compares it to a standard RNN. To ensure a fair comparison with a similar number of trainable parameters, we used a hidden size of 68 for the standard RNN. Additionally, we applied a ReLU(tanh(·)) activation function in the RNN’s hidden layer to match the activation used for the GFR neurons.

Our results demonstrate that the GFR-RNN model outperforms the standard RNN on the L-MNIST task, achieving a test accuracy of 0.958 compared to the RNN’s accuracy of 0.890. Figure Appendix C.2 illustrates the log train loss of both models over 300 training epochs, showing a faster decrease and lower convergence point for the GFR-RNN model. These findings underscore the potential of our GFR neurons to enhance RNN performance on tasks requiring memory across various time scales.

## Appendix C.2. Sequential MNIST using GFR with Biological Parameters

To create a GFR-RNN with GFR neurons with biological parameters, we randomly sampled 128 GFR neurons (Δ*t*, Δ*t*′ = 20) with explained variance ratio greater than 0.7 on Noise 2 to generate the hidden layer of the GFR-RNN.

**Table Appendix C.1:** Accuracy on the L-MNIST task.

| **Model** | **# Parameters** | **Train Accuracy** | **Test Accuracy** |
| --- | --- | --- | --- |
| RNN | 7354 | 0.898 | 0.890 |
| GFR-RNN | 7434 | 0.968 | 0.958 |

We initialized the recurrent connections by sampling from *N* (1.5, 3), as using the default initialization scheme resulted in GFR neurons getting stuck in the flat region of the activation function before the firing threshold. We multiply the input **x**_*t*_ into each GFR neuron by their respective maximum current. In addition, we normalized the input and recurrent input into each GFR neuron by the input layer dimension and hidden layer dimension respectively. We consider two GFR-RNNs: one where the GFR neuron parameters are frozen and one where they are not. Doing so, we get the following result:

**Table Appendix C.2:**
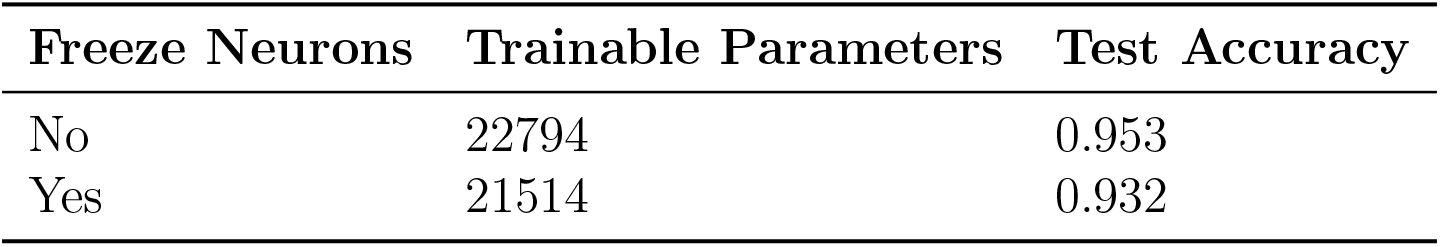
Test accuracy on L-MNIST.

| **Freeze Neurons** | **Trainable Parameters** | **Test Accuracy** |
| --- | --- | --- |
| No | 22794 | 0.953 |
| Yes | 21514 | 0.932 |

## Appendix

**D. Societal Impacts**

While this dataset does not have any clear societal implications, future biologically informed machine learning projects making use of it could, like other machine learning tools, have either positive or negative impacts on society.

## Appendix

**E. Acknowledgments**

S.M. has been in part supported by NSF 2223725, NIH R01EB029813, and RF1DA055669 grants. We also wish to thank the Allen Institute founder, Paul G. Allen, for his vision, encouragement, and support.

## Appendix

**F. Declaration of Interests**

The authors declare no competing interests.

## Appendix

**G. Declaration of Generative AI and AI-assisted Technologies**

During the preparation of this work, the author(s) used ChatGPT improve the clarity and wording of the text. After using this tool, the author(s) reviewed and edited the content as needed and take(s) full responsibility for the content of the publication.

